# Early-life scarcity adversity biases behavioral development toward a bipolar-like phenotype in mice heterozygous for CNTNAP2

**DOI:** 10.1101/2024.04.18.589746

**Authors:** Gabriele Chelini, Tommaso Fortunato-Asquini, Enrica Cerilli, Katia Monsorno, Benedetta Catena, Ginevra Matilde Dall’O’, Rosa Chiara Paolicelli, Yuri Bozzi

## Abstract

The etiological complexity of psychiatric disorders arises from the dynamic interplay between genetic and environmental vulnerabilities. Among the environmental components, early-life adversities (ELA) are a major risk-factors for developing a psychiatric disorder. Yet, the mechanistic interaction between ELA and genetic vulnerability contributing to psychopathology is poorly understood. To fill this gap, we took advantage of the ideally controlled conditions of a pre-clinical approach. In this study we raised a mouse model with genetic predisposition to multiple psychiatric disorders (autism spectrum, schizophrenia, bipolar disorder), the *Cntnap2^+/-^* mouse, with limited bedding and nesting (LBN), a well-established paradigm to induce early-life stress in rodents. These mice were compared to LBN-raised *Cntnap2^+/+^* littermates, as well as parallel groups of *Cntnap2^+/+^* and *Cntnap2^+/-^* raised in standard conditions. Using a battery for behavioral phenotyping we show that ELA shapes non-overlapping phenotypic landscapes based on genetic predisposition. Specifically, we found that LBN-raised *Cntnap2^+/-^* mice develop a perseverative risk-taking behavior in the elevated plus maze and that this behavior is highly predictive of their success in the social interaction, assessed with the 3-chamber test. This finding suggests that the intrusion of anxiety into the social behavioral domain contributes to extreme gain- or loss-of function in social interaction, resembling a bipolar-like phenotype. Finally, we show that LBN promotes synaptic hypertrophy in the basolateral nucleus of the amygdala, but only in *Cntnap2^+/-^* raised in LBN this condition was found in combination with microglia abnormalities. We conclude that the interplay between ELA and *Cntnap2* haploinsufficiency exacerbates bipolar-like behaviors in mice, and that this may be consequence of deficient synaptic homeostasis in the basolateral amygdala.

## INTRODUCTION

Early-life adversities (ELA) and genetic vulnerabilities are major risk factors for the development of psychiatric disorders (1,2). Yet, the mechanistic interaction between ELA and genetic vulnerability contributing to psychopathology is largely understudied, partially because of the unavoidable difficulty of disentangling the relative contribution of these two factors in the context of clinical and epidemiological studies. This lack of knowledge limits our capacity of understanding the neurobiological mechanisms underlying specific psychiatric traits, constraining the development of targeted therapeutic strategies and social programs to improve the quality of life of the patients. In this study we developed a “double-hit” mouse model to study the interplay between ELA and a specific genetic vulnerability in the emergence of central psychiatric traits. To do so, we used the mouse model with heterozygous deletion of the CNTNAP2 gene. In the human population, homozygous mutation of CNTNAP2 is responsible for a severe neuropsychiatric disorder characterized by autistic-like symptoms, mental retardation, severe epilepsy and cortical dysplasia (3). Knock-out mice lacking the CNTNAP2 gene (Cntnap2^-/-^) successfully recapitulate this condition (4,5). To the contrary, current literature lacks a consensus on the specific association between CNTNAP2 heterozygous mutation and an exclusive neuropsychiatric diagnosis (6–10). People carrying heterozygous CNTNAP2 mutation present a phenotypic spectrum that ranges from neurotypical to autism spectrum (ASD)(3), bipolar disorder (BD)(9,10), and schizophrenia (SZ)(9,10). Importantly, according to current literature, rodents carrying the CNTNAP2 mutation (Cntnap2^+/-^) do not display significant deviations from their *wild-type* (WT) littermates, from a behavioral standpoint (4,11). We reasoned that, the phenotypic heterogeneity related to CNTNAP2 haploinsufficiency may arise as a consequence of the gene-environment interaction, shaping specific phenotypic trajectories in response to early-life experiences occurring during critical stages of neurodevelopment. A previous study explored this possibility by exposing *Cntnap2^+/-^* to a prenatal Poly-I:C maternal immune activation, which exacerbates autistic-like behaviors in this model (12). In this work we sought to determine whether an environmental stressor during early-postnatal development may shape behavioral hallmarks associated with a specific CNTNAP2-related conditions. To do so, we raised a group of *Cntnap2^+/-^* with limited bedding and nesting (LBN), a well-established paradigm to induce early-life stress in rodents(13). WT mice raised in LBN show slight, but meaningful signs of altered stress and anxiety regulation, highly influenced by fundamental sex-differences (13–17). Interestingly, these alterations were found in combination with signs of hypertrophy and gain-of-function in cellular, molecular and functional characteristics of brain areas involved in emotion regulation such as the basolateral nucleus of the amygdala (BLA) and the prefrontal cortex (13–18).

In this study, litters composed of both *Cntnap2^+/+^* and *Cntnap2^+/-^* raised in LBN (from this point onward defined as WT-LBN or HET-LBN respectively) were compared with parallel litters of standard reared *Cntnap2^+/+^* and *Cntnap2^+/-^* (defined as WT-CTRL or HET-CTRL respectively) to investigate the specific effect of the variables: gene, environment, sex, and their interactions. Mice were evaluated with a behavioral battery to assess multiple phenotypic traits associated with psychiatric condition: stress and anxiety, social interaction, repetitive behaviors, and context-based locomotor activity. Our results show that the multiple combinations of genetic and environmental vulnerabilities give rise to alternative behavioral outcomes, specifically driving HET-LBN mice toward a BD-like phenotype, while not promoting the emergence of autistic-like traits. Finally, in the same animals, we provide evidence for synaptic hypertrophy and aberrant microglia morphology in BLA, suggesting a potential synaptic-driven mechanism governing the abnormalities in the affective behaviors resulting from the combination of CNTNAP2 haploinsufficiency and early-postnatal stress.

## MATERIALS AND METHODS

### Animals

All experimental procedures were performed in accordance with Italian and European directives (DL 26/2014, EU 63/2010) and were reviewed and approved by the University of Trento animal care committee and Italian Ministry of Health. Animals were housed in a 12h light/dark cycle with unrestricted access to food and water. All sacrifices for brain explant were performed under anesthesia and all efforts were made to minimize suffering. A total of 108 age-matched adult mice (weight 25–35 g) were used for the behavioral study. All mice were generated from our inbred CNTNAP2 colony with C57BL/6 background.

#### Breeding strategy

To minimize the effect of confounding factors on the variables of interest (i.e. early-life stress and genotype), we adopted a standardize breeding strategy. Age-matched (postnatal age P=150-160 days) WT *wild-type* females were crossed with heterozygous males, eliminating the effect of maternal age and genotype on parental care. The first litter was discarded as to exclude the consequences of first experience in maternal care. All second-born litters were included in the study, pairing a litter raised in early-life stress condition with a control litter raised in standard cages and born in close temporal proximity (< 3 days distance). (Figure 1A shows a schematic representation of the breeding strategy).

**Figure 1.**
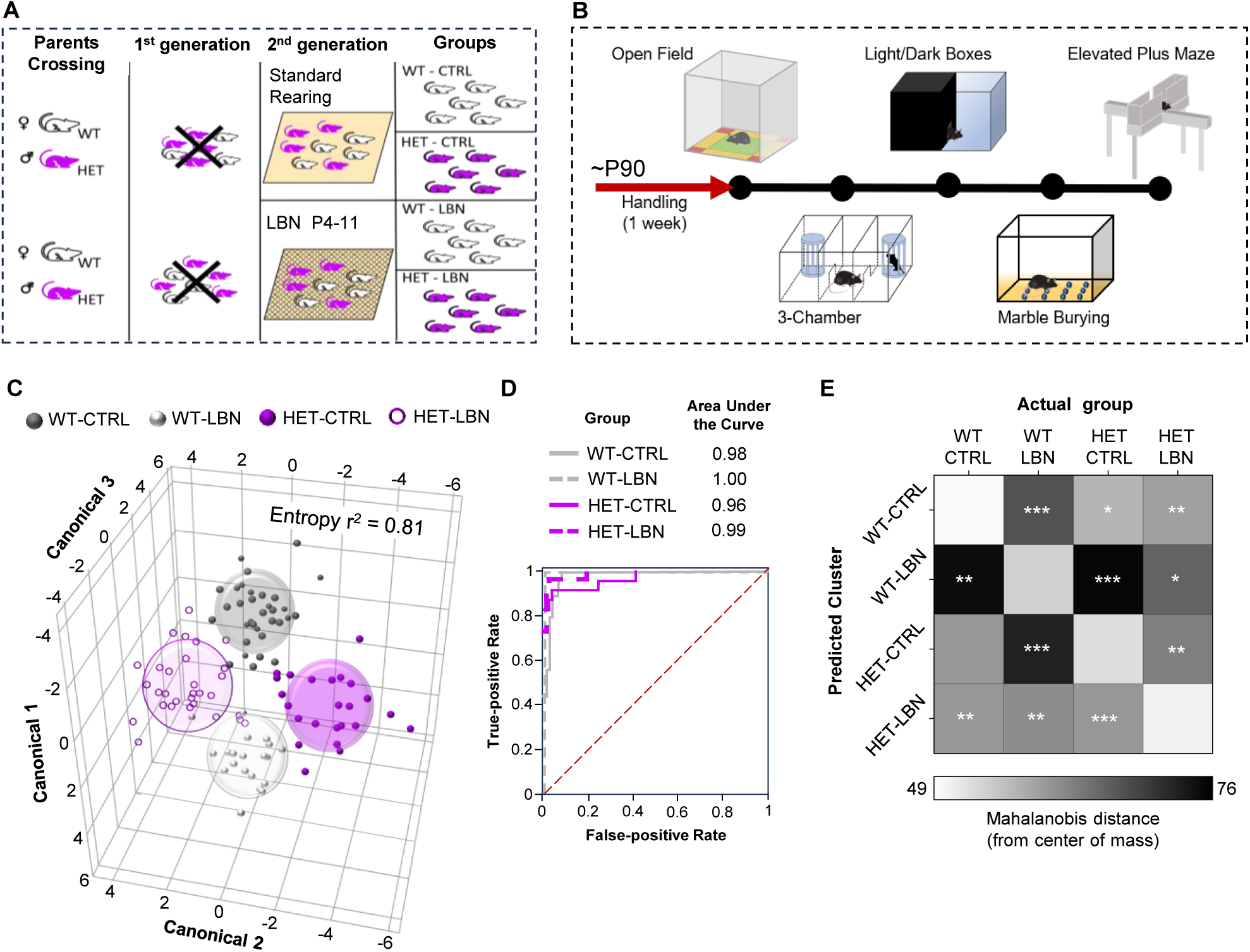
Generation and assessment of the four of experimental groups. **(A)** Schematic representation of the breeding strategy for the generation of the experimental groups. **(B)** Outline of the behavioral battery used for the assessment of stress and anxiety and autistic-like behaviors. **(C)** Discriminant analysis confirmed that the four experimental conditions give rise to four distinct behavioral phenotypes. **(D)** ROC curve, displaying nearly perfect prediction accuracy of the discriminant algorithm, confirms that the behavioral assessment successfully harnessed the phenotypic differences between the four experimental conditions. **(E)** Graphical representation of the average distance of each experimental group from the center of mass of each identified cluster. For each cluster, the average distance from its corresponding group is compared with the average distance from all other groups. *p<0.05 / **p<0.001 / ***p<0.0001.

### Limited bedding and nesting

The limited bedding and nesting paradigm was adapted from Walker et al.(13). At postnatal day 4 (P4), an aluminum mesh was placed at 2.5 cm distance from the bottom of the housing cages and the nestled material was reduced to ¼ with respect to a standard cage. The mother with the full litter was returned to a standard breeding cage once the pups reached P11.

### Behavioral battery

A specialized battery was designed including well-established test to assess both stress and anxiety domain as well as autistic-like behaviors. The battery lasted for 5 days, with each test administered daily (Figure 1B). During the week prior to the beginning of the battery, animals were daily acclimated with the room and the experimenter for five consecutive days, including 1-2 minutes daily session of gentle handling. On test days, each mouse was acclimated with the experimental room for 20 minutes before the beginning of the test. All tests were recorded by an overhead camera secured above the apparatus. For all test, apart from the marble burying, mice were tracked using the software EthoVisionXT (Noldus, Wageningen, the Netherlands). The quantification of dependent variables was automatically provided by the features included in the EthoVisionXT. Dependent variables of interest and their acquisition settings were selected prior to the beginning of the first batch of experiments, based on their relevance with respect to each test, and kept consistent across the entire study. A full list of the outcome measures used in the study is summarized in Table 1. To investigate the time-dependencies of some behavioral traits, each test (except for the marble burying) was analyzed by splitting the total time of the test (10 minutes) in two separate trials (0-5 minutes/5-10 minutes).

**Table 1.**
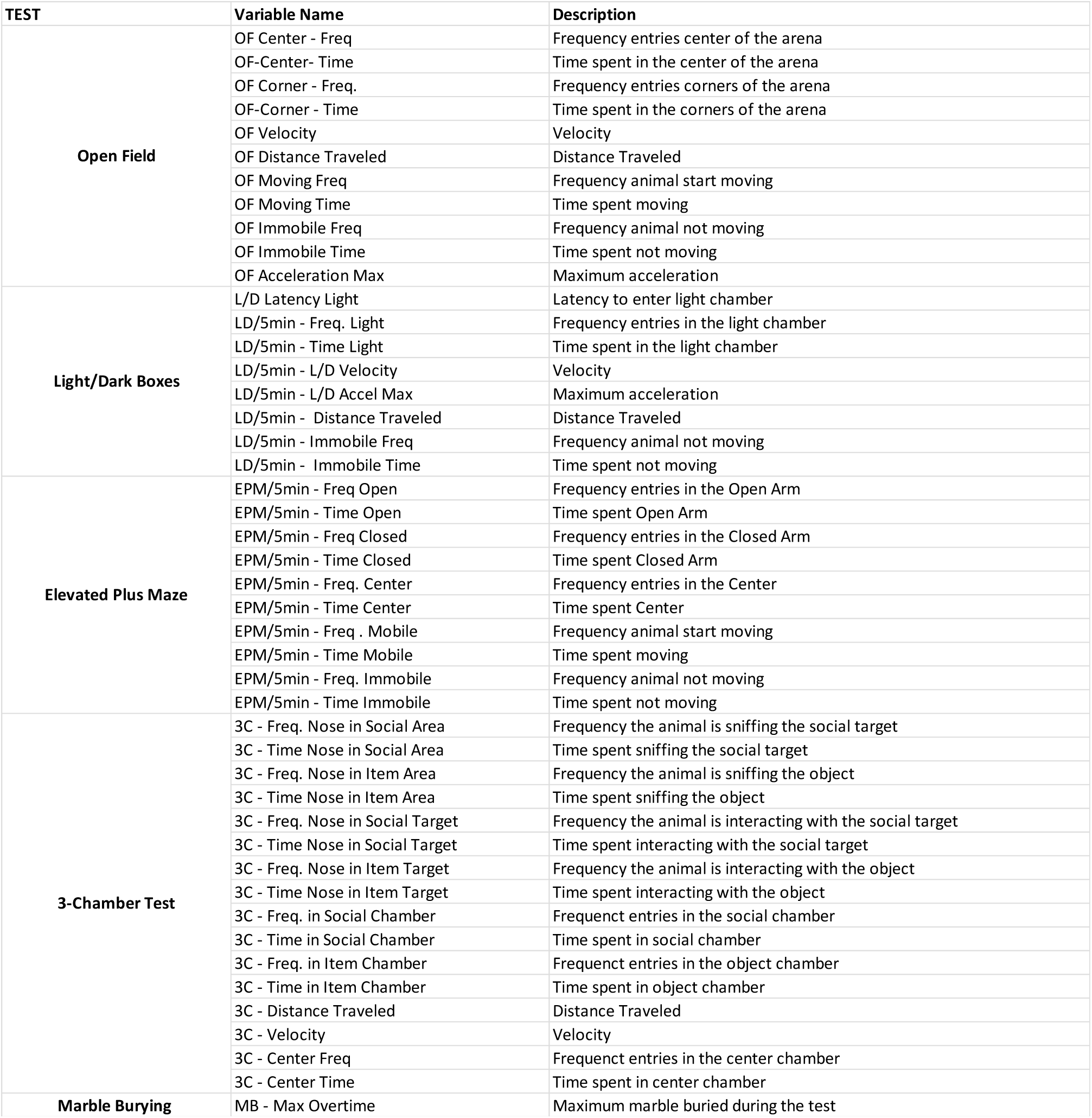
List of the outcome measures used in the behavioral assessment for each of the test. All these variables were included in the discriminant analysis.

#### Open field (OF)

Mice were placed and let free to navigate an open space in a squared arena (40 × 40 × 40 cm) for a total of 20 minutes while vide-recorded(19). Only the results of the first 10-minutes are reported in this work as to match the timeframe of the other stress and anxiety tests.

#### Three-chamber test

The apparatus consisted of a plexiglass rectangular box (60 × 40 × 22 cm), each chamber (20 × 40 × 22 cm) having blacked colored external walls and transparent panels separating the 3 chambers (20). Immediately after the open field test, happening always the day before the 3-chamber, mice were placed in the three-chamber apparatus and allowed to freely explore it for 20 min. On test day, mice were put in the apparatus and let free to navigate for the first 10 minutes. Then, the sociability test started by introducing one unfamiliar mouse placed into a wire cylindrical cage (20cm in height, 10 cm bottom-diameter, 1cm bars spaced) in one of the corners of one of the lateral chambers. An identical empty wire cage was placed in the equivalent corner of the opposite chamber. The test mouse was allowed to freely interact with the unfamiliar mouse or the empty cage for 10 minutes. No differences were found between the first and the second half of the 3-chamber test, thus we only present here the results for the total time of the test (10 minutes).

#### Ligh/Dark boxes (L/D)

Mice we introduced into an arena (40×25×25) split in two halves (chambers) separated by a black panel (21). One half of the arena had the walls externally coated in black, fully shielding it from the light. The other half had transparent walls and a bright light source positioned nearby. A 5×8 door allowed the mice to freely transition between the two halves. Animals were allowed to freely move into the apparatus for a total of 10 minutes while videotaped.

#### Marble Burying

Five cm of standard mouse bedding were put in a standard rat cage (26 cm × 48 cm × 20 cm) and 16 standard glass toy marble were position in 4 rows equidistant from each other (22). Mice were let free to navigate the cage for 30 minutes while videotaped. For every 5 minutes of the test, a snapshot of the video was taken to quantify the number of marbles buried at that interval. 1 point was assigned to marble covered for more than 75% of their visible surface; 0.5 points were assigned when a marble was buried for less than 75%. No major informative differences were found in this test; thus, the results are only reported in the supplementary Figure 1.

#### Elevated Plus Maze (EPM)

Mice were place at the center of a standard elevated plus maze apparatus, consisting of a two open arms (25 × 5 × 0.5 cm) across from each other and perpendicular to two closed arms (25 × 5 × 16 cm) and a center platform (5 × 5 × 0.5 cm) (23). Animals were allowed to freely move into the apparatus for a total of 10 minutes while videotaped.

### Definition of indexes used in the behavioral assessment

In each formula ‘T’ stands for “time”, ‘F’ stands for “frequency”. *OF*. To assess the subject preference for a safe zone (corners), versus an anxiogenic one (center of the arena) we used an index defined as: *Safety Preference Index = T Corners / (T Corners + T Center).* The preference for arena borders against the center (thigmotaxis) was defined as: *Thigmotaxis = T Borders / (T Borders + T Center)*.

#### EPM

To assess the subject preference for a safe exploration (center of the maze) versus an anxiogenic ones (open arms) we used two indexes defined as: *Safe Exploration Index = F Center / (F Center + F Open Arms); Safe Exploration Index = T Center / (T Center + T Open Arms).* The preference for safe areas (closed arms) versus anxiogenic ones (open arms) was defined as: *Safety Preference Index = F Closed Arms / (F Closed Arms + F Open Arms); Safety Preference Index = T Closed Arms / (T Closed Arms + Time Open Arms)*

For all tests, the calculated indexes span from 0 to 1, where 1 represents complete preference for a safe choice while 0 represent complete preference for the risky choice. The social preference was calculated using the formula: *Social Preference Index = T sniffing the mouse / (T sniffing the mouse + T sniffing the empty object)*.

### Tissue processing and immunofluorescence

One month after the end of the behavioral battery, mice were deeply anesthetized with isoflurane and sacrificed by decapitation. Brains were excised and split in two halves. The right hemisphere was freshly dissected in multiple brain regions, snap frozen in dry ice and preserved at -80° for future studies. The left hemisphere was washed in 0.1% phosphate buffer saline (PBS) and post-fixed overnight in 4% paraformaldehyde (PFA), switched to a cryoprotectant solution (80% PBS, 20% glycerol with 0.1% sodium azide) and stored at 4°C. Cryoprotected brains were sectioned on a vibratome (Leica, VT1200) at 40μm thickness. Serial sections were collected in 24 separate compartments and stored at 4°C in cryoprotectant solution.

#### Synapse study

Free-floating slices were rinsed three times in PBS (10 min each), then washed in PBS containing 0.2% detergent (Triton-X, Fisher, AC215680010) for 30 minutes. Tissue sections were then incubated in blocking solution [2% Bovine serum albumin (BSA), 1% fetal bovine serum (FBS) in PBS] for 3 hr and then transferred to primary antibody solution [2% BSA, 1% FBS, 1 to 1000 dilution of primary antibody (Rabbit anti-PSD95, Abcam, ab18258)] and incubated at room temperature for 24 hr. Then, sections were rinsed three times in PBS (5 min each) and placed in a fluorophore-conjugated (Alexa Fluor^TM^ Plus 488) secondary antibody solution [1:500 dilution of donkey anti-Rabbit secondary antibody (Thermo Fisher, AB_2762833) in PBS] for 24 hr. Sections were then washed 5 min in PB, mounted on superfrost slides, dried for 1 hr and coverslipped with fluorescent mounting medium (Southern biotech 0100-01).

#### Microglia study

Floating slices were permeabilized in 0.5% Triton X-100 in PBS for 90min at RT, followed by incubation with blocking buffer (2% BSA in permeabilization buffer) for 1h at RT. Primary antibodies were diluted in blocking buffer (Iba1, 1:1000, Wako cat no 019-19741; CD68, 1:400, Bio-Rad cat no MCA1957) and incubated overnight at 4°C with mild agitation. Brain slices were then washed in PBS at RT for 3 times, 10 minutes per wash, and further incubated with Alexa Fluor Plus-labelled secondary antibodies (1:1000 in blocking buffer) for 2h at RT. After additional 3 washes in PBS, nuclei were stained for 10 minutes at RT with DAPI (1μg/ml in PBS). Finally, slices were mounted on microscope slides using Mowiol 4-88 (Sigma Aldrich, cat no 81381). For both studies, slides were stored at 4°C in the dark until use.

### Confocal microscopy and image analysis

*Synapse study*. A confocal laser scanning microscope Leica TCS-SP8, equipped with a HC PL APO 40x objective and interfaced with Leica “LAS-X” software was used. Images were recorded at a resolution of 1024 pixels square, 400 Hz scan speed with a zoom factor of x5 to maximize puncta visualization. Excitation/emission wavelengths were: 490/520 for Alexa-488 fluorophore. Acquisition parameters were set during the first acquisition and kept consistent for all the images. For each subject, six size-matched (36×36 μm) z-stack images were acquired from the BLA using a pseudo-random strategy, caring to exclude cell bodies from the images and focusing exclusively on the dendritic neuropil. Each stack spanned for a total of 10 μm thickness, with a step-size of 0.5 μm. The maximum intensity projection for each z-stack was post-processed using the “subtract background” and “despeckle” features within the imageJ software (NIH, USA)(24) and quantified with the “3D object counter” plug-in. Postprocessing and detection settings were kept consistent across all images.

#### Microglia study

Acquisitions were taken at a Stellaris 5 confocal microscope (Leica), with a 63x objective and digital zoom of 1 (Z-stack, step size 0.3μm). Identical imaging settings and resolution (1024×1024 pixels) were maintained across all the tested conditions for each experiment. Microglia morphometry and engulfment analyses were carried out using the Imaris software (Bitplane). 3D reconstruction was performed with the built-in “Surface” function by applying the same thresholds across all the tested conditions. All entire microglia cells per field of view were reconstructed and analyzed.

### Statistical analysis

All statistics were computed using the software JMP-pro17. The discriminant analysis algorithm included all the relevant metrics obtained in the behavioral assessment (Table 1). For the behavioral studies, a 3×3 multiple regression model was applied to test for the main effects of covariates *early-life experience* (ELE), *genotype* (Gen) and *sex*, and their interactions. Whenever the variable sex was found not significant by the regression model it was excluded from the statistical analysis. When the variable sex was found to significantly impact the results, but in equivalent degree for all 4 experimental, a multiple regression model was applied to exclude the effect of this covariate from the data. No major effect of the interaction between sex and other variables was found in most of the studies, thus the results of multiple comparisons reported in the main text do not report sex differences. The only test showing a meaningful sex-bias was the L/D boxes, for which we report the sex-dependent multiple comparisons in Supplementary Figure 3. Multiple comparisons were calculated using Tuckey’s correction. To compare the first half of each test (0-5 minutes) with the second half (from 5 to 10 minutes) we used a 2-way mixed model ANOVA; multiple comparisons were calculated using Sidak’s correction. An extensive report of the statistical analysis is reported in the Supplementary file n.1.

## RESULTS

### Early-life adversity determines distinct behavioral phenotypes based on genetic background

First, we sought to determine whether the combination of gene and environment may result in major, non-overlapping, phenotypic domain. In the behavioral assessment we looked for both autistic-like behaviors and signs of stress and anxiety (Figure 1A). After the completion of the behavioral battery, we performed a discriminant analysis using the four experimental conditions (WT-CTRL / WT-LBN / HET-CTRL / HET-LBN) as predictors and the outcome measures of all behavioral tests as descriptors (Table 1, Figure 1A, 1B). The discriminant algorithm successfully determined the existence of four distinct clusters (Figure 1C) which corresponded, with virtually perfect accuracy (Figure 1D), to the four experimental conditions. Using the mahalanobis distances as a measure of proximity of each subject from the center of mass of every cluster, we confirmed that all experimental groups are significantly closer to their predicted cluster than any of the others (Figure 1E). The only exception to this was the HET-CTRL cluster, which resulted not significantly apart from the WT-CTRL group (Figure 1E). This observation is in line with previous studies where no major differences were found between *Cntnap^+/+^*and *Cntnap2^+/-^.* Our findings establish that the LBN approach determines multidimensional behavioral landscapes based on genetic predisposition, giving rise to four distinct phenotypic categories to investigate.

### CNTNAP2 haploinsufficiency mitigates ELA-derived anxiogenic behavior in the open field

As a first step in the behavioral assessment, we looked for signs of generalized anxiety in the open field (Figure 2A). At the 0-5 minutes interval, we highlighted a main effect of LBN, which resulted in a significant increase in the safety preference index in both WT- and HET-LBN groups compared to WT-CTRL (Figure 2B). This was accompanied by a significant increase in the time spent in the corners specific for the WT-LBN group (Figure 2C). No group difference was found in the 5-10 minutes interval, while the main effect of LBN persisted (Figure 2D). These findings suggest that early exposure to scarcity-adversity is sufficient to exacerbate an anxiogenic phenotype, but this effect is partially mitigated by *Cntnap2* haploinsufficiency. Additionally, we used multiple motility descriptors to assess potential discrepancies in the locomotor activity. We found that none of the parameters analyzed (velocity, distance traveled, time and frequency mobile and immobile) were affected by the experimental conditions (Figure 2 E, F and supplementary Figure 2), as none of the groups diverged from the WT-CTRL. This establishes that neither haploinsufficiency of C*ntnap2* nor the LBN negatively impact motor abilities nor exacerbate hyperactive behavior in mice.

**Figure 2.**
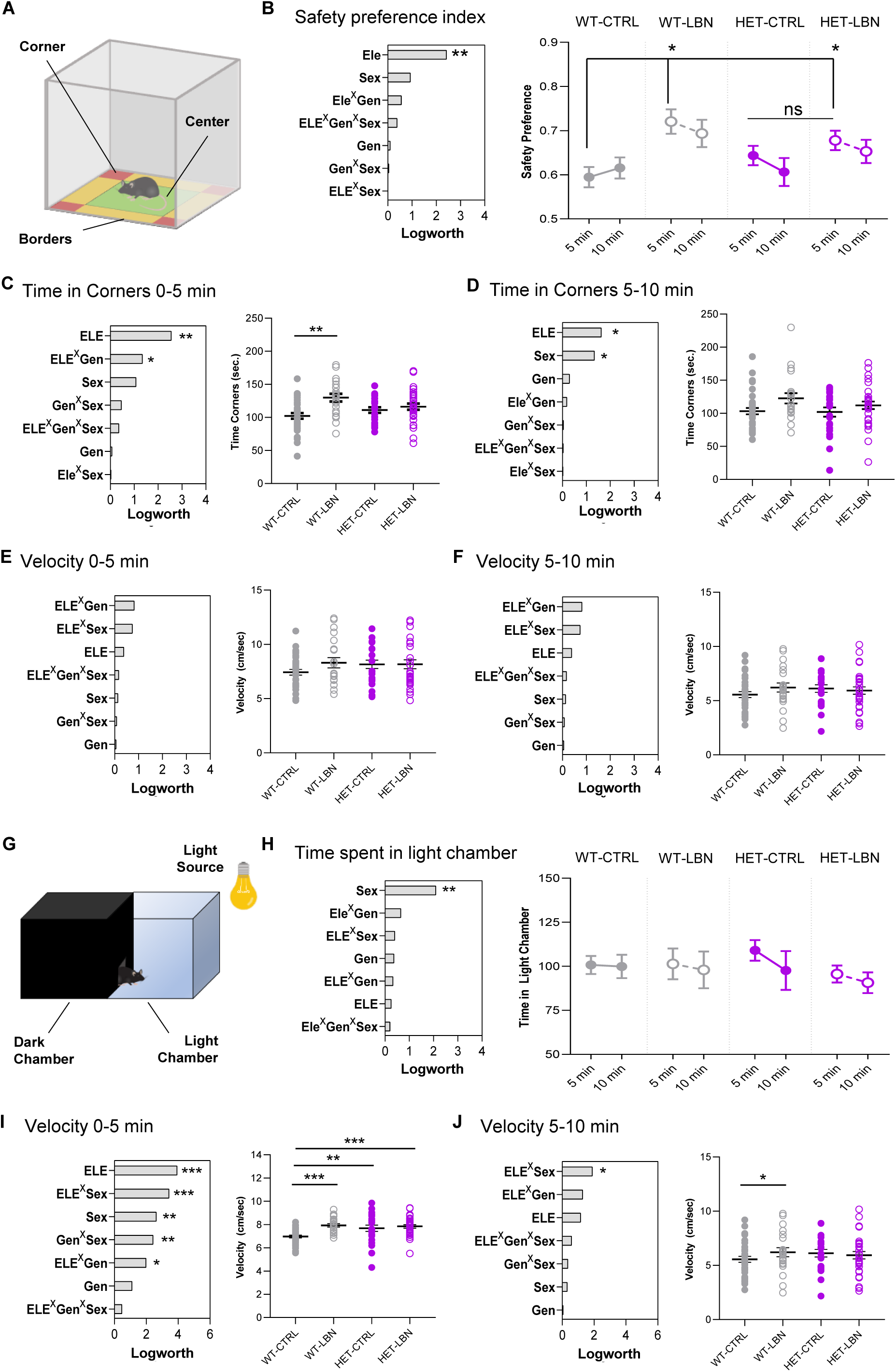
ELA induces anxiogenic behaviors that are mitigated by Cntnap2 haploinsufficiency. **(A)** Graphical depiction of the Open Field arena and its partitions. **(B-F)** Left panels: charts summarizing the quantification of the main effect of the covariates included in the model; Right charts: depict the results of multiple comparisons for the given outcome measure. **(B)** A main effect of ELE determines a preference for the safe areas of the OF arena (corners) in both WT and HET-LBN groups, specifically during the first 5 minutes of the test. **(C)** During the first 5 minutes of the test, WT-LBN spend significantly more time in the corners compared to WT-CTRL, highlighting a main effect of both early-life experience (ELE) and the interaction between ELE and genotype (ELExGen). **(D)** A main effect of the ELE persists during the 5-10 minutes interval, but group differences are abolished. **(E, F)** No differences in the velocity indicates that neither ELS nor CNTNAP2 mutation have an impact on the locomotor activity. **(G)** Graphical depiction of the L/D arena and its partitions. **(H)** A remarkable effect of sex was found in the time spent in the light chamber, but no impact of either ELA or Genotype**. (I)** Both ELE and Genotype, as well as sex have a significant impact on the mice velocity in the light chamber during the first 5 minutes of the L/D test, resulting in a significant increase in all groups compared to WT-CTRL. **(J)** Only WT-LBN group maintain significantly higher speed in the second half of the test. See Figure S2 for further details on sex differences. Abbreviations: ELE= early-life experience; Gen= genotype; ^X^ indicates interaction. *p<0.05 / **p<0.001 / ***p<0.0001.

### Both LBN and CNTNAP heterozygous mutation independently result in a hyperactive phenotype in the L/D transition test

To further explore the effect of ELA onto anxiogenic behavior, we challenged the four experimental groups in the L/D test, evaluating their risk-assessment strategy in a condition of perceived danger (i.e. light chamber, Figure 2G). In this setting, no differences were found between the four experimental groups in the time spent in the light chamber, at neither time-point (Figure 2H). However, during the first 5 minutes only, we observed a main effect of the variable sex that was primarily driven by an increase in the time spent in light displayed by the males of the WT-LBN group, compared to females (Supplementary Figure 3A). This sex-bias was abolished in the HET-LBN (Supplementary Figure 3B). Consistently, WT-LBN males showed to cover more distance during their exploratory activity in the light chamber (Supplementary Figure 3 C, D). This finding confirms a previous work showing that LBN defines male-specific behavioral deviations in WT animals (25). However, by looking at the locomotor activity, we found a striking effect of both LBN and genotype onto the velocity within the light chamber, showing that all 3 experimental groups are statistically different from WT-CTRL (Figure 2I) during the first 5 minutes of the test. Only the WT-LBN group showed a protracted change in the 5-10 minutes interval (Figure 2L). Altogether these findings suggest that, when mice are provided with the option of exploring an aversive environment, both ELA and *Cntnap2* haploinsufficiency determines a severe hyper-active behavior, but only WT-males express a drastic change in their risk-assessment strategies. This confirms the results of the OF, showing that *Cntnap2* heterozygous mutation partially prevent the negative consequence of early-postnatal stress.

### Early-life stress exacerbates perseverative risk-taking behavior in *Cntnap2^+/-^* mice

To further understand potential changes in risk-assessment strategies, we tested the four groups in the elevated plus maze (EPM). In this test we evaluated the mice tendency to risk-taking (walking in the open arm) against their natural predisposition to explore the surroundings from the center of the maze as an indicator of innate curiosity (Figure 3A). We found that WT-CTRL mice display a drastic increase in the safe exploration index between the first and second half of the test, privileging a safe choice over a dangerous one after an initial risk-assessment. This trend is preserved in both WT-LBN and HET-CTRL mice (although to a lesser degree in this latter group), but completely abolished in HET-LBN mice, highlighting a main effect of the gene × environment interaction (Figure 3B and Supplementary Figure 4B). These data are determined by an increase in both the frequency (Figure 3C, D) and the total time (Figure S4 C, D) HET-LBN mice spend in the open arms of the maze (Figure 3E) exclusively during the 5-10 minutes time-window.

**Figure 3.**
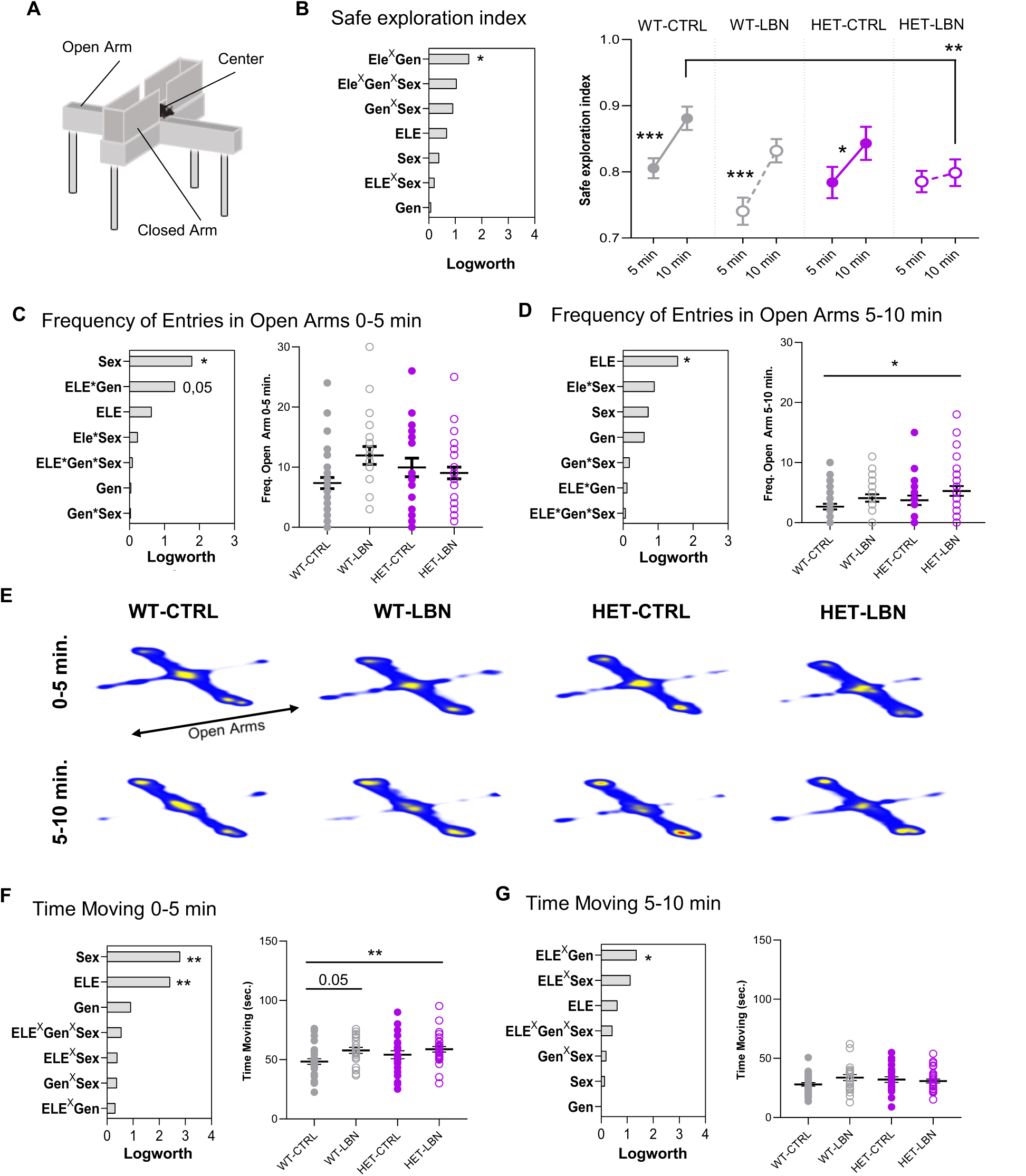
HET-LBN mice display a perseverative risk-taking behavior in the EPM. **(A)** Graphical depiction of the elevated plus maze and its partitions. **(B-D, F, G)** Left panels summarize the quantification of the main effect of the covariates included in the model. Right charts depict the results of multiple comparisons for the given outcome measure. **(B)** A main effect of the interaction ELE^X^Gen affects the mice preference for a safe area (center) versus a dangerous one (open arms); during the second half of the test, the safe exploration index increases in WT-CTRL, WT-LBN and HET-CTRL, but not in HET-LBN, resulting in a significant difference with WT-CTRL. **(C)** During the first 5 minutes, the frequency of entries into the open arm does not change in neither of the groups and only a shared effect of Sex is significantly explaining the model. **(D)** During the 5-10 minutes interval, a main effect of the variable ELE is explained by a significant increase in the frequency of entries in the open arm specific for the HET-LBN group compared to WT-CTRL. **(E)** Representative heatmaps depicting mice activity at both the time-points of the test; note how LBN exposure to the open arm persists beyond the first 5 minutes. **(F)** A main effect of LBN was found in the time moving in the maze for the 0-5 minutes interval, due to a slight increase in motility in WT-LBN and a large increase in HET-LBN. **(G)** Group differences in the motility state are abolished in the second half of the test, while a small effect of the interaction ELE^X^Gen emerge. Abbreviations: ELE= early-life experience; Gen= genotype; ^X^ indicates interaction. *p<0.05

This finding suggests that, irrespectively from their early-life experience, mice possess an innate ability of tuning their emotion-driven behavior to privilege a safe choice against a potentially dangerous alternative. This ability is dampened by *Cntnap2* haploinsufficiency and exacerbates in a perseverative risk-taking behavior when *Cntnap2^+/-^* mice are raised in LBN. Finally, we discovered a main effect of LBN on the locomotor activity only during the first 5 minutes (Figure 3 F, G), with a subtle increase affecting both WT-LBN and HET-LBN mice, which corroborates our previous finding in the L/D test.

### Successful social interaction in LBN-raised *Cntnap2^+/-^* correlates with their risk-taking behavior, suggesting an alternative social-behavioral pattern

To test for potential effects in the social behavioral domain, we assessed the social preference of the four groups in the 3-chamber test (Figure 4A). At first, no main effect of variables sex, early-life experience or genotype, nor group differences could be appreciated, at neither time points (Figure 4B). Indeed, all groups showed similar time spent sniffing the social target (Figure 4C) as well as successful social preference (index above 0.5, Figure 4D). Then, we asked whether any of the changes in anxiety regulation might play a role in the outcome of the social interaction. To explore this concept, we correlated the time spent sniffing the social target with all the measures extrapolated in the stress and anxiety evaluation (Figure 4E). In the HET-LBN group, we found that both the time and frequency of exposure to the open arm during the second half of the EPM were highly predictive of the time the mice invested exploring the co-specific (Figure 4E). Thus, we hypothesized that, for *Cntnap2^+/-^* mice raised in LBN, successful social interaction might be contingent on their willingness to take a risk. To verify this hypothesis on the basis of our data, we calculated a regression model including the variable “sniffing time” and “frequency open arms” (Figure 4F). Then we used the slope coefficient of the regression to correct the values of the “sniffing time” after excluding the effect of preservative risk-taking from the variance. Doing so, we found a main effect of the variable LBN in reducing the sniffing time, primarily derived by a significant reduction affecting the HET-LBN group compared to both WT- and HET-CTRL (Figure 4G). This reduction resulted in the loss of the social preference in the HET-LBN group selectively (Figure 4H). These findings suggest that an intrusion of anxiety into the social behavioral domain may alter the social interaction pattern in HET-LBN mice. To test this hypothesis, we used a principal component analysis (PCA) including all the measures related to the social interaction obtained in the three-chamber test (see supplementary file 1 for a detailed list). Doing so, we identified a total of five significant principal components (PC), three of which sufficient to explain 90% of the variance (Figure 4I). Using this three leading PC, we performed an iterative unsupervised *k-mean* clustering. Using the elbow method (Figure 4I) (26), we determined the presence of four distinct modalities of social interaction (Figure 4I), that we named sociophenotypes (SPs). Using the social preference index as an indicator, we categorized the four SPs as hypersocial (SP1), asocial (SP2), mildly social (SP3) and normo-social (SP4) (Figure 4J and Supplementary Figure 5A). Finally, analyzing the frequency distribution of the four SPs within the four experimental groups we found a significant difference between the WT-CTRL and the HET-LBN, resulting from a disproportionate prevalence of the SP3 in the HET-LBN group (Figure 4K). These data were further confirmed by multiple correspondence analysis showing that HET-LBN and SP3 segregate together, and in isolation from all the other categories (Supplementary Figure 5 B). We conclude that, while not impaired in their social-behavioral skills, HET-LBN mice are more likely to express extreme gain or loss of function in their levels of social interaction, with a larger portion of subjects displaying lack of social preference, and that this condition appears to be dependent on their affective state.

**Figure 4.**
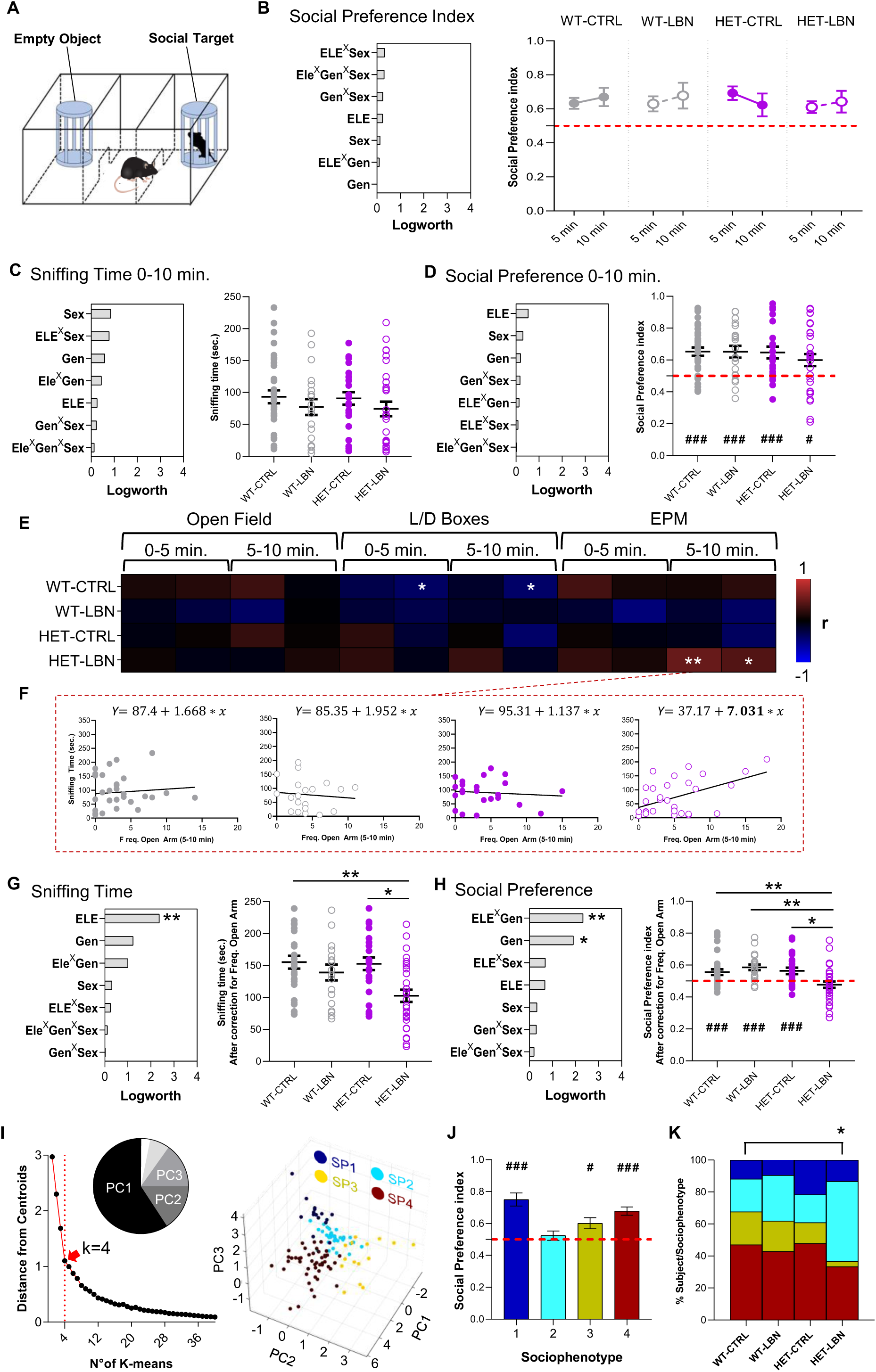
Altered social-behavioral pattern in LBN-raised *Cntnap2^+/-^* mice. **(A)** Graphical depiction of the Open Field arena and its partitions. **(B-D, G, H)** Left panels summarize the quantification of the main effect of the covariates included in the model. Right charts depict the results of multiple comparisons for the given outcome measure. **(B)** No changes in the social preference at neither time-point. **(C, D)** Neither the time of social interaction **(C)** nor the social preference **(D)** are impacted by experimental conditions. **(E)** Correlation matrix between the time spent sniffing the social target and the outcome measures obtained in the stress and anxiety assessment. **(F)** Regression plot showing the linear relationship between the time spent sniffing the social target and the frequency of entries into the open arm. **(G, H)** Results of the 3-chamber test after correcting the data for the effect of risk-taking behavior: **(G)** A main effect of the variable ELE results in HET-LBN mice spending less time sniffing the social target compared to both WT- and HET-CTRLs; **(H)** By consequence, HET-LBN display a lack of social preference due to a main effect of the ELE*Gen interaction. **(I)** Iterative k-mean clustering using the elbow method determined the presence of four distinct sociophenotypes (SPs); the pie chart shows the relative weight of each PC in explaining the variance; the scatter plot displays the elbow method to select the optimal number of *k* for the clustering; the 3D scatter plot depicts the separation of the four SPs. **(J)** Differences in the social preference across SPs; only SP2 shows significantly impaired social behavior. **(K)** Graphical depiction of the frequency distribution (%) of the SPs within each of the four experimental groups; note that SP2 is over-represented in the HET-LBN group, making it significantly divergent from WT-CTRL. Abbreviations: ELE = early-life experience; Gen = genotype; ^X^ indicates interaction. *p<0.05 / **p<0.001 / ***p<0.0001 / # indicates social preference significantly different by chance: 0.5

### Early-life stress forces excitatory synapses hypertrophy in the BLA

Then we hypothesized that the behavioral rigidity displayed by HET-LBN mice in the EPM may be linked to deficient synaptic turnover in a brain area dedicated to emotional processing. To investigate this possibility, we assessed the density and the size of PSD95-positive punctate in the BLA (Figure 5A). We observed a close-to-significance effect of the variable LBN in both parameters (Figure 5B, C). To get a better understanding of the impact of ELA onto synaptic phenotype we analyzed the frequency distribution of the sizes of all the PSD95 puncta size identified. We found that, within each genotype, there is a significant shift of the distribution toward an increase in the percentage of larger post-synaptic densities (Figure 5 D, E). Thus, we sorted the PSD95-puncta into three size-based categories (small: from 30 to 70 voxels, medium: from 71 to 110 voxels and large from 111 to 150 voxels) and calculated the ratio between the number of puncta of each category against both of the others (Figure 5F-H). We found no difference in the ratio between small-to-medium sized puncta (Figure 5F). Conversely, a significant effect of LBN reduced both the small-to-large as well as the medium-to-large ratios (Figure 5G, H). Additionally, the medium-to-large ratio was significantly decrease in the HET-LBN group compared to HET-CTRL (Figure 5H). Finally, we discovered that the size of PSD95 puncta significantly and exclusively correlates with the safety preference index in the second half of the EPM (Figure 5I). Specifically, the percentage of smaller puncta positively correlates with a preference for safety (Figure 5J) while the percentage of medium and larger puncta associates with reduced preference for safety (Figure 5K, L). Altogether these findings suggest that ELA promote a decrease in the number of smaller synapses in favor of larger ones. This phenotype is worsened in *Cntnap2^+/-^*mice raised in LBN and may contribute to their maladaptive risk-taking behavior observed in the EPM.

**Figure 5.**
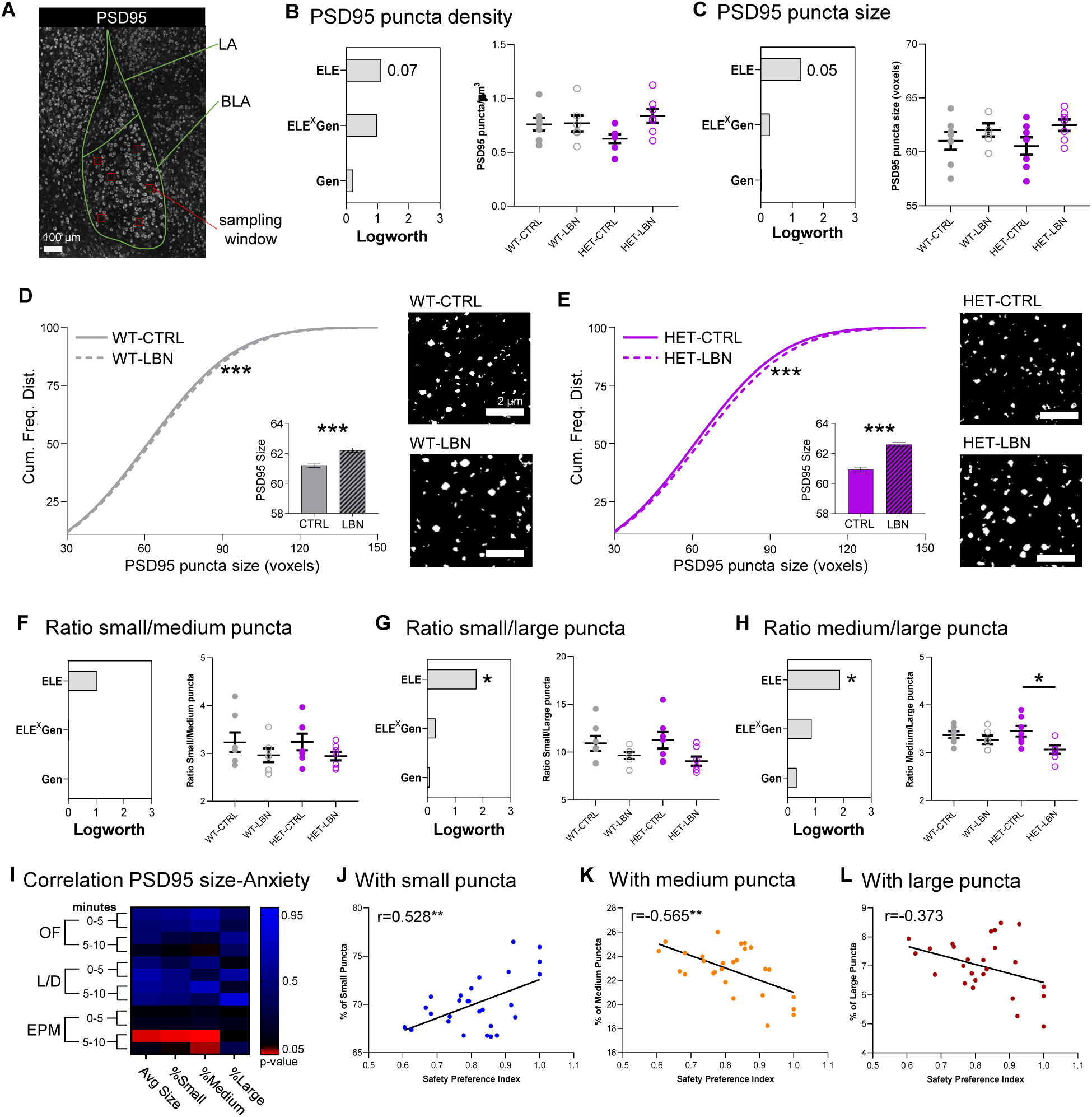
ELA promotes synaptic hypertrophy in BLA. **(A)** Confocal photomicrograph of the basolateral amygdala (BLA) showing the location of sampling windows used for the analysis of PSD95 puncta. **(B, C, F-H)** Left panels summarize the quantification of the main effect of the covariates included in the model. Right charts depict the results of multiple comparisons for the given outcome measure. **(B, C)** A close-to-significant effect of ELE was found in both the density **(B)** and size **(C)** of PSD95 puncta. **(D, E)** Analysis of the relative frequency distribution of the puncta size revealed a significant shift in favor of larger puncta in both WT-LBN and HET-LBN groups compared to their respective WT-CTRL and HET-CTRL. **(F)** Ratio between small and medium puncta size is unaffected by the experimental conditions. **(G)** The ratio between small and large puncta is significantly affected by ELE, but does not result in group differences. **(H)** The ratio between medium and large puncta is significantly affected by ELE, with a stronger effect displayed in HET-LBN, significantly diverging from HET-CTRL. **(I)** The size of PSD95-puncta selectively correlates with the safety preference index during the 5-10 minutes interval of the EPM. **(J)** The percentage of smaller-sized puncta positively correlates with the safety preference while **(K)** the percentage of medium-sized puncta is negatively correlated. **(L)** Weak negative correlation between the percentage of larger spines and safety preference (p=0.05). Abbreviations: ELE= early-life experience; Gen= genotype; ^X^ indicates interaction. *p<0.05 / **p<0.001 / ***p<0.0001.

### *Cntnap2* haploinsufficiency determines a vulnerability state in microglia that exacerbate in a hypertrophic phenotype after ELA

We then asked whether synaptic alterations in the BLA may be associated with microglia abnormalities. Microglia are emerging as key players in brain pathologies, including neurodevelopmental and neuropsychiatric disorders (27–29). Notably, microglial impairments are often associated with abnormal synapse density and function, leading to behavioral alterations in sociability, cognition and anxiety(30–32). We stained for IBA1 and CD68 to visualize and analyze microglial morphology and phagolysosomes, respectively (Figure 6A, B). Morphometric analysis on three-dimensional cell reconstruction, revealed that WT-LBN and HET-CTRL microglia did not present any alteration compared to WT-CTRL, whereas HET-LBN microglia displayed almost a two-fold increase in volume (Figure 6B-D). Consistently, we also found that the number and total volume occupied by CD68+ structures were unchanged in WT-LBN and HET-CTRL microglia compared to WT-CTRL, but significantly higher in HET-LBN (Figure 6B, E, Supplementary Figure 6A). Conversely, the average volume of phagolysosomes was increased in HET mice, with a significant genotype effect (Figure 6F). Next, we asked whether the higher content of CD68 particles per cell in HET-LBN could be just a mere consequence of microglial hypertrophy. When normalizing the number and volume of CD68 particles taking into account the microglial volume, we observed an increased CD68 content only in HET-CTRL, while this effect was absent in HET-LBN (Supplementary Figure 6B, C).

**Figure 6.**
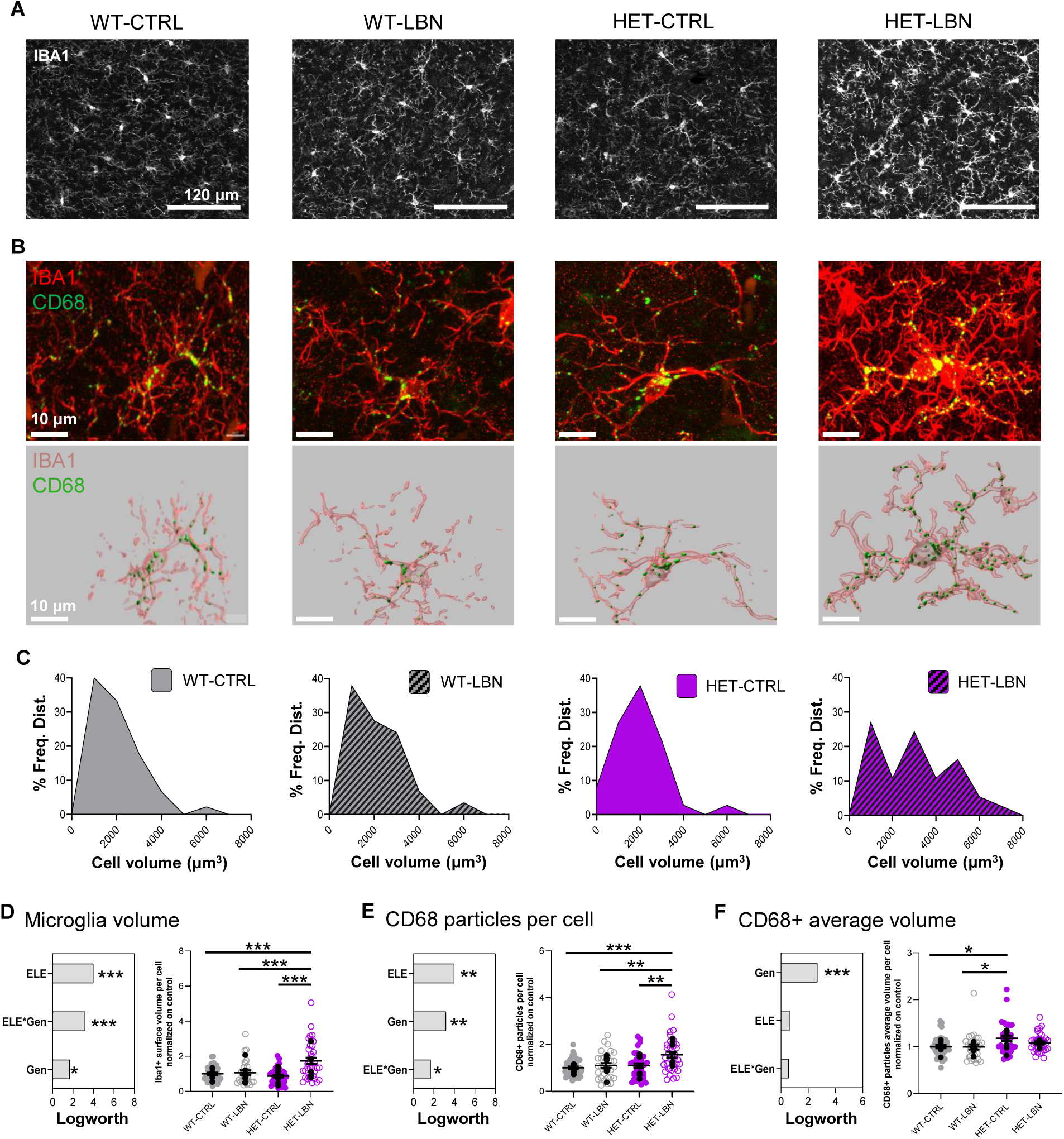
Altered microglia phenotype in Cntnap2^+/-^ mice is worsened by ELA. **(A)** Confocal photomicrographs showing an overview of IBA1-positive microglia cells in BLA. **(B)** Confocal photomicrographs capturing a detailed view of a typical microglia appearance in each of the four experimental groups (top) and paired with its relative 3D reconstruction (bottom). **(C)** Frequency plots showing the distribution of microglia size in the four experimental conditions as revealed by rank analysis; HET-LBN presents with a greater proportion of larger cells (between 4000 and 6000 μm^3^) compared to WT-CTRL and HET-CTRL (for visualization purpose, cells were categorized according to size-bins of 1000 μm^3^). **(D, E)** Left panels: charts summarizing the quantification of the main effect of the covariates included in the model; Right charts: depict the results of multiple comparisons for the given outcome measure. **(D)** A main effect of both ELE and ELE^x^Gen is driven by significant increase in the microglia size selective for HET-LBN (as forecast in panel C). **(E)** The number of CD68+ phagolysosomes per cell is selectively increased in HET-LBN, determining significant effect of both ELE and Gen and their interaction. **(F)** Conversely, the average volume of CD68-positive phagolysosomes per cell surface area is exclusively affected by CNTNAP2 haploinsufficiency, with significant differences observed between HET-CTRL and both WT groups. Black dots indicate values for single animal. Colored dots indicate values of single cells. Abbreviations: ELE= early-life experience; Gen= genotype; ^X^ indicates interaction. *p<0.05 / **p<0.001 / ***p<0.0001.

Overall, these data suggest that the *Cntnap2* haploinsufficiency alone is not sufficient to affect microglial morphology, but that exposure to ELA is required to observe alterations in these cells. On the other hand, the phagolysosomal compartment is affected in control *Cntnap2* haploinsufficient mice, and the interaction with ELA dampen these alterations.

## DISCUSSION

In this study we combined a genetic vulnerability to neuropsychiatric disorders (*Cntnap2* heterozygosis) with an early-life stressful event to investigate the interplay between environmental stress and genetic predisposition in the emergence of transdiagnostic psychiatric hallmarks. First, we established that ELA biases behavioral trajectories based on genetic predisposition, by showing that a discriminant component algorithm can assign each subject to its respective experimental condition based on the results of the behavioral assessment, with nearly perfect accuracy. Then, looking at the specific outcome of each test, we discovered that exclusively the *Cntnap2^+/-^* mice raised in LBN express a purposeless risk-taking behavior by persistently walking into the open arm of the elevated plus maze through the entire duration of the test. We further observed that HET-LBN animals are more likely to show higher levels of social avoidance, and that their engagement in social exploration may be dependent on their excessive risk-taking tendencies. We conclude that altered stress and anxiety regulation biases social-behavioral strategies in HET-LBN selectively, determining an ambivalent patter of hyper- or hypo-sociability. Translationally, these alterations resemble those of people diagnosed with bipolar disorders, as they show extreme positive or negative levels of both anxiety and social behavior and a manic-like risk taking tendency in a potentially harmful environment.

HET-LBN mice seem to lack the ability of fine-tuning their exploratory behavior when exposed to a potential threat (falling from the open arm of the EPM), suggesting a maladaptive context-dependent emotional cognition. We hypothesized that this sign of behavioral rigidity may be accompanied by deficient synaptic flexibility in the BLA. This brain region works as a multisensory hub, integrating interoceptive and exteroceptive inputs to encode the biological value of external cues, based on adaptive needs (33). Using the size of PSD95 puncta as a proxy for synaptic strength, we found that ELA promote the accumulation of larger postsynaptic densities in the BLA at the expense of smaller ones, and that this effect is amplified by *Cntnap2* heterozygous mutation. Elegant studies show that the fine-tuning of receptive fields (34) or the specificity of associative memories (35) benefit of a rebound synaptic downscaling that restores local network dynamics to a critical state (36). A highly dynamic synaptic turnover is an ideal feature that allows the maintenance of this homeostasis in local brain circuits (34–40). We postulate that a similar mechanism may be at play in this model. Lack of synaptic turnover within the BLA may underlie a deteriorated signal-to-noise ratio within the amygdala microcircuitry, determining an over-generalization of behavioral strategies irrespectively to the biological valence of the surrounding environment. In support of this hypothesis, a previous study show that organized synaptic ensembles identified by immunolabeling of 6-sulfated chondroitin sulfate (41) are reduced in the postmortem brain of people with BD (42), suggesting that the lack of locally organized plasticity may contribute to BD pathology. In line with this hypothesis, we interrogated the microglia phenotype within the BLA of the same mice showing altered synaptic phenotype. Microglia are major players implicated in activity-dependent synaptic refinement and studies are increasingly showing the contribution of this cell-type in the pathophysiology of psychiatric disorders (28,29). Furthermore microglial morphological and functional alterations have been frequently observed across mouse models of neurodevelopmental disorders (43) (Hughes, Moreno, Ashwood 2023), including *Cntnap2* full knockouts mice (44) (Dawson et al. 2023). Our findings show that microglia are affected by *Cntnap2* heterozygous mutation, particularly when challenged with ELA. HET-LBN mice display bigger microglial volume and higher content of CD68-positive phagolysosomes within the BLA compared to all the other conditions. Altered CD68 content can be associated with increased or decreased phagocytosis of synapses and cellular debris (31,32,45). However, here it remains to be determined whether this modulation is functionally involved in the refinement of synapses and in the behavioral alterations described in this study.

Importantly, we confirmed that standard-reared *Cntnap2*^+/-^ mice do not show major behavioral abnormalities; however, we were able to detect some sub-threshold discrepancies with respect to WT-CTRL (locomotor activity in the L/D, attenuated adaptation in the EPM). Furthermore, both the synaptic and the microglia phenotypes in HET-CTRL showed some critical differences with WT-CTRL, suggesting that a divergent neural functioning may underlie parallel behavioral strategy that eventually converge to similar behavioral outcomes. Interestingly, similar evidence come from the study of family members of patients with a diagnosis of BD (46); these studies show that undiagnosed first-degree relatives display functional (47), molecular (48) and behavioral (49) characteristics overlapping with what observed in their diagnosed family member. In lines with this finding, we propose that C*ntnap2* haploinsufficiency may confer a status of subliminal emotional vulnerability that exacerbates in maladaptive behaviors in response to an early-life adverse experience.

Finally, we would like to emphasize a few methodological considerations. One is the importance of using a standardized breeding strategy. This approach is already suggested in previous articles describing the LBN procedure(13). In this work, however, it was necessary to also controlling for the effect of maternal genotype. As revealed by our in-depth behavioral phenotyping, *Cntnap2^+/-^* females do show minimal discrepancies with controls that went previously undetected. Thus, there was some chance that the LBN may interact with the maternal genotype differently, preventing us to establish whether behavioral alterations of the offspring are due to the offspring or the mother genetic background; the use of *wild-type* dams eliminated this problem. A second critical point is the inclusion of the factor time in the behavioral evaluation. The time-dependency of symptoms is an integral component of psychopathology(50,51). Especially in mood disorders, symptoms cycles on multiple time-scales, from seasonal(52), to circadian(53) up to milliseconds differences in reaction times(54) and discrepancies in time-perception(55). In this work we chose to investigate arbitrary time-bins to obtain a behavioral redout of characteristics such as impulsivity or stereotypies, which proved an effective strategy to harness the complexity of some deviant traits. It will be necessary, in future studies, to expand the assessment of time-behavior interaction in preclinical model of psychiatric disorders.

A major limitation of this study is that it primarily focused on the behavioral characterization, and only marginally digs into the neurobiological underpinnings. The synaptic and microglia anomalies in the BLA are -most likely-just the tip of the iceberg of a broader scale impact of LBN onto *Cntnap2^+/-^.* Additional studies are required to elucidate which other brain systems are affected, how does the gene-environment interaction influence local and broad-scale connectivity; whether certain types of neurons are impacted and to what extent. As many questions remain open, we emphasize the fact that, in this study, we leveraged a not-fully-penetrant genetic vulnerability to develop a functionally relevant preclinical model of psychiatric pathology, delivering a timely application of a negative postnatal experience. These results carry a threefold advantage. First, they provide the scientific community with an improved ecological model for the study of BD-like phenotypes in laboratory mice. Second, they offer a framework that could be applied to multiple other animal models with genetic vulnerability to deepen our understanding of the dynamic interplay between genetic and environmental predispositions in the pathophysiology of mental disorders. Finally, they shed some light on the phenotypic heterogeneity associated with C*ntnap2* heterozygous mutation, suggesting that mutations of this gene may constitute a causal factor for specific psychiatric pathology only and exclusively when associated with specific environmental factors, such as prenatal immune activation (12) or early-life stress.

## Supporting information

Supplementary Material - statistic report

## Acknowledgments

We thank all the administrative and technical staff of CIMeC for support. A special thank goes to Mrs. Michela Maffei, animal caretaker of the CIMeC animal facility, for her endless care and support provided in managing the animal colony used in the study and Dr. Tommaso Pecchia for helping with the technical implementations of some of the behavioral apparatuses. GC effort was covered by the ‘CARITRO postdoctoral fellowship’, funded by Fondazione Cassa di Risparmio di Trento e Rovereto and the University of Trento starting grant, provided by the University of Trento. YB was supported by the Strategic Project TRAIN - Trentino Autism Initiative from the University of Trento (grant 2018-2022). RCP was supported by grants from ERC (StGrant REMIND 804949), SNSF (SNSF 310030_197940) and funding from UNIL.

## Author Contributions

**GC** conceptualized the work, designed and performed all experiments, analyzed the data, provided funding and wrote the manuscript. **TFA** contributed to establish the mouse colony for the early-life stress and the behavioral experiments. **EC, BC, GMD** contribute to the research work. **KM** carried out the microglia analysis and wrote the manuscript paragraph for the microglia study. **RCP** supervised the microglia and synapses studies and edited the manuscript. **YB** supervised the study, provided funding, and edited the manuscript.

