## Supplementary Material - Supporting Data and Figures for "Early-life scarcity adversity biases behavioral development toward a bipolar-like phenotype in mice heterozygous for CNTNAP2"

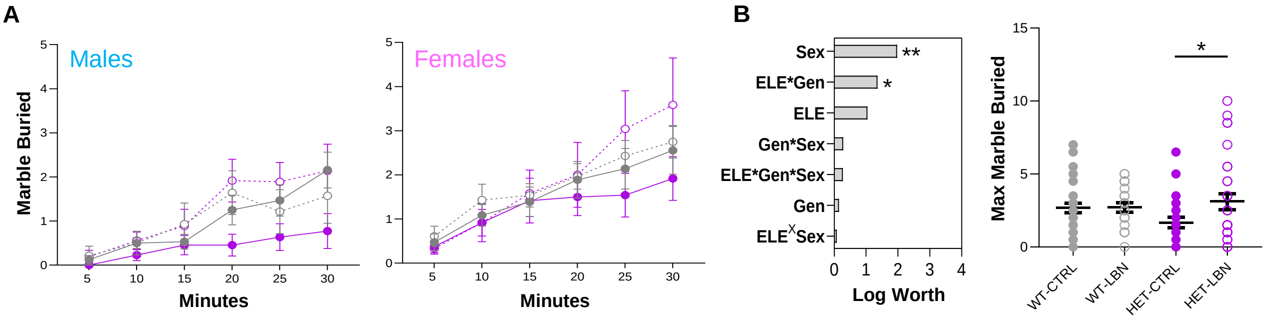


**Supplementary Figure 1. No major changes in motor stereotypies were found in the marble burying.**
**(A)** Across the multiple time-points, females show a trend of burying more marble than males. **(B)** Left charts summarize the quantification of the main effect of the covariates included in the model. Right charts depict the multiple comparisons for the maximum number of marble buried during the whole duration of the test. After correction for the effect of the variable 'sex' we found a main effect of the ELE*Gen interaction due to a significant increase in the marble buried by HET-LBN mice compared to HET-CTRL. Neither of these group, however, diverged from WT-CTRL. These data may indicate that a minor decrease in digging behavior due to *Cntnap2* haploinsufficiency is normalized after ELA. Abbreviations: ELE= early-life experience; Gen= genotype; $^{X}$ indicates interaction. $^{*}$p<0.05 / $^{**}$p<0.001

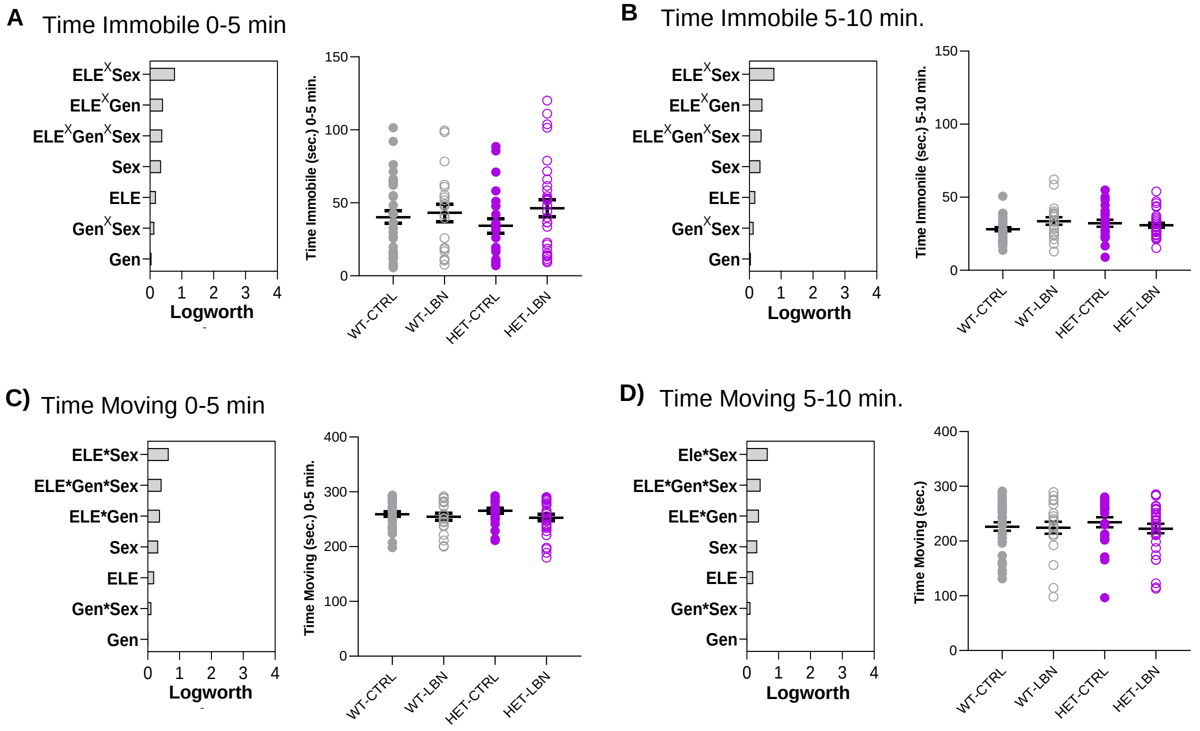


**Supplementary Figure 2. Locomotor activity in the OF is unaffected by ELA and genotype**
For each panel **(A-D)** Left charts summarize the quantification of the main effect of the covariates included in the model. Right charts depict the multiple comparisons for the given outcome measure. For either time-points, no main effect of the covariates, nor group differences were detected in the time spent immobile **(A, B)** nor the time spent moving **(C, D).** Abbreviations: ELE= early-life experience; Gen= genotype; $^{X}$ indicates interaction.

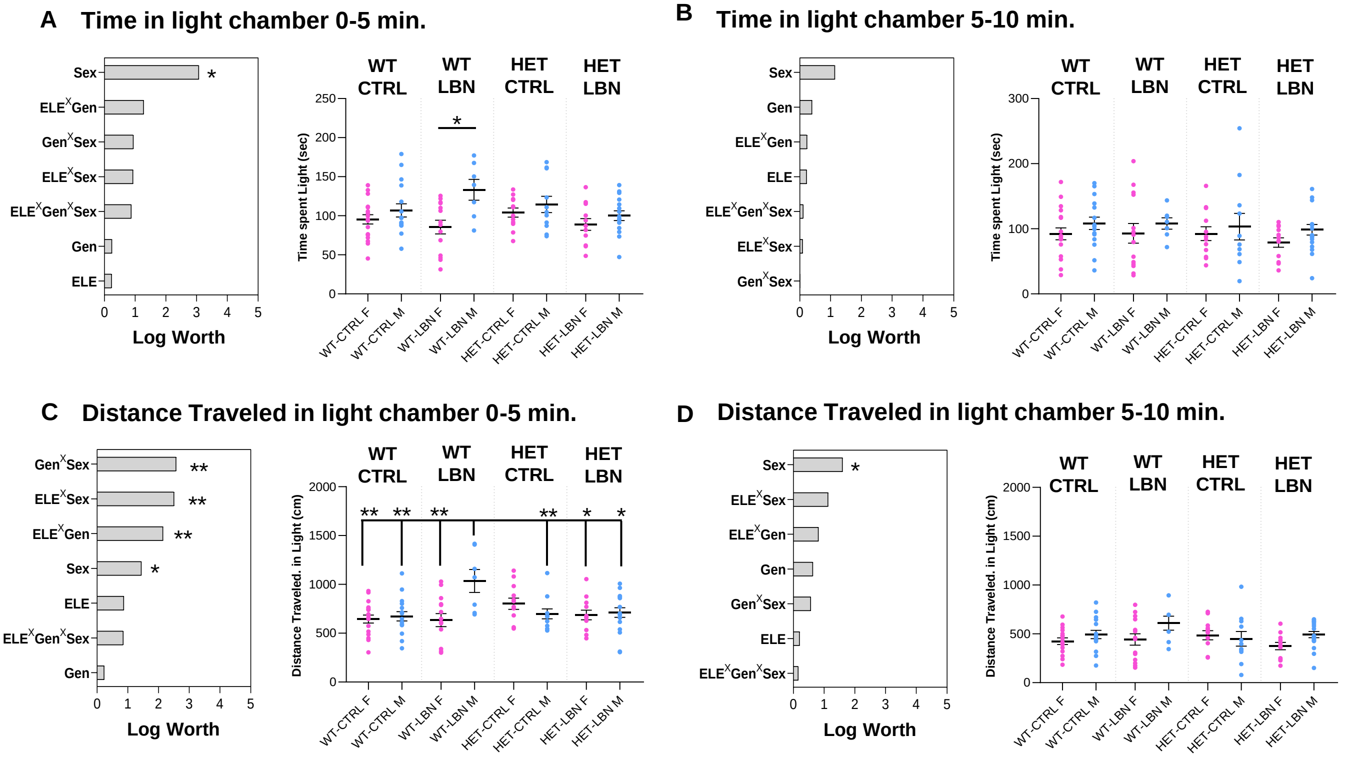


**Supplementary Figure 3. Sex-differences in the L/D boxes**
**(A-D)** Left panels summarize the quantification of the main effect of the covariates included in the model. Right charts depict the results of multiple comparisons for the given outcome measure. **(A)** During the first 5 minutes, a significant effect of sex indicates that males spend more time in the light chamber compared to group-matched females; this effect is significantly enhanced within the WT-LBN group**. (B)** All Differences are abolished in the 5-10 minutes interval**. (C)** Consistently with (A), WT-LBN males cover more distance exploring the light chamber compared to other groups, except for HET-CTRL females. **(D)** All Differences are abolished in the 5-10 minutes interval, only a marginal effect of sex persists. Abbreviations: ELE= early-life experience; Gen= genotype; $^{X}$ indicates interaction. $^{*}$p<0.05 / $^{**}$p<0.001.

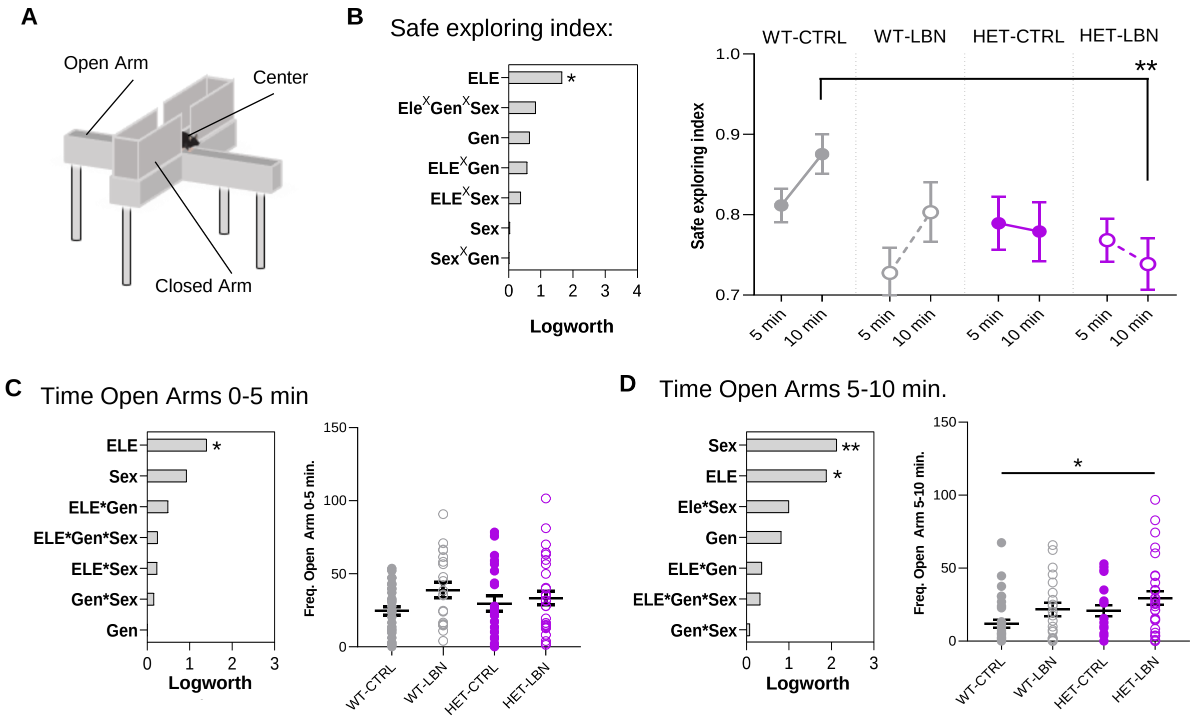


**Supplementary Figure 4. Results of the EPM using the time open arm as outcome measure closely match the frequency data**

**(A)** Graphical depiction of the Elevated plus maze and its partitions **B-D)** Left panels summarize the quantification of the main effect of the covariates included in the model. Right charts depict the results of multiple comparisons for the given outcome measure. **(B)** A main effect of the variable ELE drives the HET-LBN group to a preference for the open arms in the second half of the test. **(C)** During the first 5 minutes, a main effect of ELE does not result in significant group differences. **(D)** During the 5-10 minutes interval, a main effect of the variable ELE is explained by a significant increase in the time spent in the open arm displayed by the HET-LBN group compared to WT-CTRL. Abbreviations: ELE= early-life experience; Gen= genotype; $^{X}$ indicates interaction. $^{*}p<0.05$ / $^{**}p<0.001$

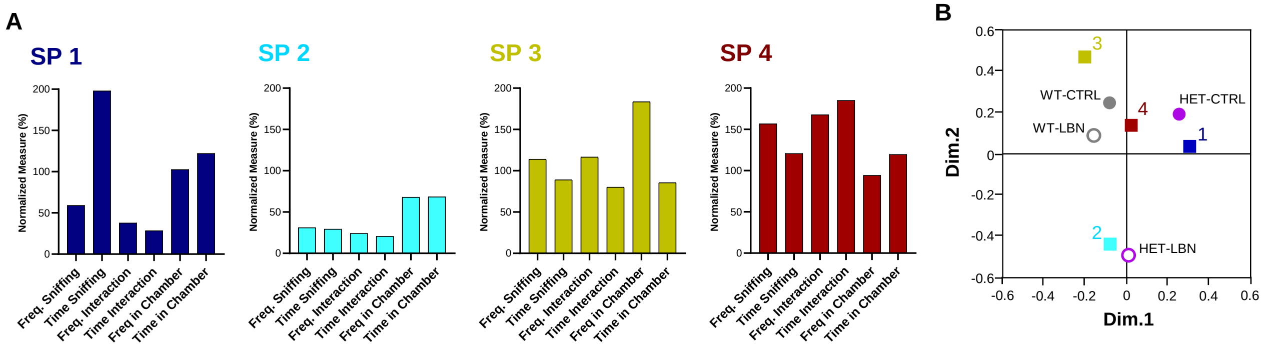


**Supplementary Figure 5. Altered social-behavioral pattern in Cntnap2$^{+/-}$ raised in LBN**
**(A)** Qualitative characterization of the four SPs based on the original metrics included in the PCA. To obtain comparable results across different measurement, the values obtained from each subject is expressed as a percentage of the group average. SP1 is characterized by extreme levels of social curiosity, with lower expression of direct social interaction. SP2 show low level of all the social-behavioral descriptors. SP3 emerged as an intermediate category, with average results in all descriptors. SP4 showed high, but balanced, levels of all social components. **(B)** Multiple correspondence analysis displaying the similarities of the four SPs with the four experimental groups confirms the association between SP2 and HET-LBN.

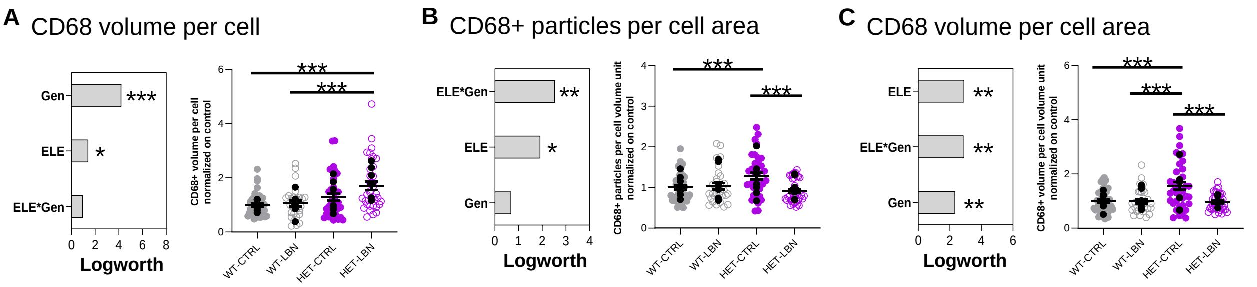


**Supplementary Figure 6. CD68 phagolysosomes are highly impacted by Cntnap2 heterozygous mutation**
**A-C) Left panels:** charts summarizing the quantification of the main effect of the covariates included in the model; **Right charts:** depict the results of multiple comparisons for the given outcome measure.
**(A)** The CD68+ volume per cell is primarily affected by CNTNAP2 mutation and significantly increased in HET-LBN compared to both WT groups. **(B)** The number CD68+ phagolysosomes per cell surface area is primarily affected by ELE*Gen interaction and significantly increased in HET-CTRL compared to WT-CTRL and HET-LBN. **(C)** The total volume of CD68-positive phagolysosomes per cell surface area is equally affected by all covariates, with significant differences observed between HET-CTRL and all other groups.
Black dots indicate values for single animal. Colored dots indicate values of single cells.
Abbreviations: ELE= early-life experience; Gen= genotype; $^{X}$ indicates interaction. $^{*}p<0.05$ / $^{**}p<0.001$ / $^{***}p<0.0001$.
